# Extracellular vesicles mediate OxLDL-induced stromal cell proliferation in Benign Prostatic Hyperplasia

**DOI:** 10.1101/2024.05.31.596872

**Authors:** Franco F. Roldán Gallardo, Daniel E. Martinez Piñerez, Kevin F. Reinarz Torrado, Gabriela A. Berg, Vanina G. Da Ros, Manuel López Seoane, Cristina A. Maldonado, Amado A. Quintar

## Abstract

2

**Background:** Clinical and basic research evidence has suggested a possible linkage of Benign Prostatic Hyperplasia (BPH) to proatherogenic conditions such as dyslipedemia and hypercholesterolemia, but the underlying mechanisms remain still unknown. We here aimed to explore the impact of dyslipidemic contexts on prostatic stromal cell proliferation and on the release of extracellular vesicles (EVs).

**Methods:** Mice were exposed to a high-fat diet and human prostatic stromal cells (HPSC) subjected to oxidized-LDL (OxLDL). Cell proliferation assays and EV characterization were performed to elucidate the involvement of EVs in the BPH.

**Results:** Pro-atherogenic conditions significantly induced proliferation in murine prostatic cells and HPSC, while metformin demonstrated a mitigating effect on OxLDL-induced proliferation. Additionally, OxLDL augmented EV production and release by HPSC, thereby promoting further proliferation, highlighting a potential mechanism underlying BPH progression.

**Conclusions:** The findings suggest that pro-atherogenic conditions contributes to prostatic cell proliferation and EV production, influencing BPH progression. Metformin emerges as a promising therapeutic avenue for BPH management. This study underscores the intricate interplay between dyslipidemia, cell proliferation, and therapeutic targets in BPH pathogenesis.

## 3 Introduction

Benign Prostatic Hyperplasia (BPH) is a chronic and progressive condition associated to uncontrolled cell proliferation in the transition zone of the prostate, which leads to the gland enlargement, manifested as lower urinary tract symptoms [LUTS^1^]. The prevalence is stimated to be around 50% in men over the age of 50, increasing by 10% for each decade of life^2^. While several pharmacological therapeutic strategies are currently available, for most patients, surgical procedures to remove excess prostatic tissue become inevitable^3^.

Changes in lifestyle and dietary habits, along with the rise of the “Western Diet” have sparked concerns regarding the impact of dyslipidemia on organ pathophysiology^4^. In fact, dyslipidemia and hyperlipidemic stages constitute the primary risk factor for atherosclerosis and cardiovascular disease^5,6^. Although it has been suggested a potential pathogenic role of such a context in the abnormal growth of the prostate gland^7,8^, the precise cellular and molecular mechanisms underlying dyslipidemia-induced proliferation remain unclear.

OxLDL, a modified form of low-density lipoprotein (LDL), undergoes oxidation, altering both its structure and functionality^9^, and plays a crucial role in initiating atherosclerotic lesions and vascular pathologies. Additionally, it triggers cellular proliferation in several tissues^10,11^. For instance, OxLDL promotes endothelial cell proliferation by activating the Rho/Akt signaling pathway^11^ and modulates cell cycle in vascular cells by regulating p27Kip1 via Rho^12^. Furthermore, it regulates proliferation, cell survival, and cell cycle in smooth muscle cells^13,14^. In the prostate, OxLDL induces the secretion of proinflammatory interleukins, potentially impacting both inflammatory and proliferative processes^14^.

Extracellular Vesicles (EVs) have emerged as crucial mediatiors of intercellular communication in various physiological and pathological conditions due to the different biomolecules they transport^15^. For instance, EVs from mesenchymal stem cells stimulate the proliferation of tubular epithelial cells^16^, and they have also been implicated in proliferation and migration of chronic myeloid leukemia cells^17^, migration and invasion^18^, and apoptosis^19^, through the transport of specific microRNAs.

OxLDL has been shown to promote EVs release by endothelial cells, regulating inflammation during atherosclerosis through miR-155^20^. Macrophage-derived EVs stimulated by OxLDL participate in cell migration in the atherosclerotic lesion^21^, while OxLDL-dependent platelet-derived microvesicles trigger procoagulant effects and amplify oxidative stress^22^. Given the diverse functions attributed to EVs in cellular communication and the impact of OxLDL on prostate cells^23^, further research is needed to elucidate the presence and functions of EVs in the hyperproliferative state of BPH. Therefore, our study aims to investigate the roles of dyslipedemia, OxLDL, and EVs in BPH by assessing their influence on mouse prostates and on human prostatic stromal cells (HPSC) derived from BPH patients.

## 4 Material and Methods

### Mice and high-fat diet (HFD)

Male C57BL/6 mice, aged 4 weeks, were housed at IBYME-CONICET and INICSA-CONICET (Argentina) and fed either standard chow diet (6% fat, 24% protein, 40.5% carbohydrates, 3210 kcal/kg, n=4) or a HFD (30.3% fat, 17.8% protein, 30% carbohydrates, 4640 kcal/kg) for 12 weeks^24^. The mice were kept under controlled photoperiod (14h light/10h darkness) with *ad libitum* access to water and food. Procedures adhered to NIH Guidelines and were approved by the local ethical committes (CICUAL, UNC, Argentina). At the end of protocol, blood was collected for total cholesterol deteermination; prostatic complexes were extracted, fixed with 4% formaldehyde, embedded in paraffin and sectioned into 4µm histological sections for further analyses.

### Treatments in primary cultures of prostatic stromal cells and THP-1 cell line

Prostate tissue samples were collected from BPH-diagnosed patients aged 50 to 70 years via TURP and placed in MCDB131 medium (Sigma-Aldrich) on ice. Mechanical and enzymatic digestion using type IA collagenase (200 U/mL, Sigma-Aldrich) was performed for 1.5 h. Cells were then seeded into 6-well culture plates in MCDB131 medium, supplemented with 15% fetal bovine serum (Internegocios S.A.). Cultures were maintained for a minimum of 15 days and underwent at least 6 passages to ensure stromal purity before experimental protocols^25^. The study was approved by the Bioethics Committee of Sanatorio Allende (Córdoba, Argentina).

Prior to OxLDL stimulation, HPSC were kept in serum-free medium for 24h. To this, LDL was obtained from human plasma and underwent oxidation with Cu2+ as previously described^26^. Two types of OxLDL samples were obtained: moderately oxidized (OxLDL1) and highly oxidized (OxLDL2). OxLDL1 was achieved by exposing LDL to Cu2+ for 4 hours at 37°C, followed by the addition of 2 mmol/L EDTA to halt the process. For OxLDL2, exposure to Cu2+ for 4 hours was conducted without subsequent EDTA addition^26^. HPSC were stimulated with OxLDL, at concentrations of 5, 20, and 100 µg/mL, metformin (2 and 10mM), atorvastatin (20µM and 2.5µM), or a combination with both for 24 h. The human monocytic THP-1 cell line (from ATCC) was cultured in RPMI1640 medium (Thermo Fisher Scientific Inc.) supplemented with 10% FBS and stimulated with OxLDL1 and OxLDL2 at a concentration of 20 µg/mL.

### Proliferation assays

Cell proliferation and viability were assessed using BrdU incorporation assay, Ki-67 immunocytochemistry, and resazurin test. Briefly, cells were seeded on coverslips in 24-well culture plates, with 10µM 5-Bromo-2’-Deoxyuridine (Invitrogen) being added for the last 6 h of treatments. Cells and prostatic tissues were fixed in 4% formaldehyde and subjected to antigen retrieval using 0.1M citrate buffer, permeabilization using 0.25% Triton X-100. Coverslips were then incubated with 1U/µL DNase (1/10, Zymo Research) diluted in Krebs buffer (Sigma-Aldrich) and subsequentenly with primary antibodies (1/30 Ki-67 Cat. 550609 and 1/200 Anti BrdU Cat. 555627, both in 1% BSA, BD Pharmingen) overnight. An Alexa594-conjugated anti-mouse antibody (Invitrogen) was used during 1 h for BrdU detection and DAPI for nuclear staining. For Ki-67, Avidin-Biotin peroxidase Complex (ABC, Thermo Fisher Scientific Inc.) was added and immunoreactivity visualized by using DAB (Sigma-Aldrich). In both techniques, 1000 cells were counted to determine the percentage of proliferating cells. For THP-1 cells growing in suspension, proliferation was evaluated by counting the total number cells using a Neubauer Chamber. Cell viability and metabolic activity of both HPSC and THP-1 cells were assessed by adding Resazurin (1µg/mL, Thermo Fisher Scientific Inc.) to treatments in 96-well plates. Absorbance was then quantified at 560nm using a GloMax® Multi Detection System spectrophotometer (Promega).

### Isolation of HPSC-derived EVs

HPSC were cultured in 100 mm diameter adherent plates; upon reaching 80% confluence, the culture medium was replaced with serum-free MCDB131 for a 24-h period followed by 20µg/mL OxLDL1 treatment for 24h. The medium was then replaced with serum-free MCDB131 for another 24-h period to allow the acumulation of EVs in the medium. Supernatants were then collected and underwent ultracentrifugation to isolate EVs according to a standardized protocol^27^. Briefly, conditioned media were subjected to 10 min at 200g for 3 cycles (G146D centrifuge, Gelec®) and 20 min at 2000g (Allegra 64R Centrifuge, Beckman Coulter) to remove cellular debris and apoptotic bodies respectively. Finally, ultracentrifugation was performed at 150,000g for 50 min (Optima MAX-TL ultracentrifuge, Beckman Coulter) to obtain a pool of EVs comprising both microvesicles and exosomes. The EVs were then resuspended in 15µL of 1X PBS and stored at -80°C until further analysis.

### Cell and EVs characterization by Transmission Electron Microscopy (TEM)

Cells were fixed with Karnovsky solution for 15 min. A post-fixation step was performed using 1% osmium tetroxide for 1h and then, samples were dehydratated in 50%, 70%, 90%, and 100% acetone for 15 min at each step. Afterwards, they were embedded in EPON Araldita resin (Electron Microscopy Sciences), mixture and allowed to polymerize at 60°C for 48 h. Thin sections (90 nm) were prepared using a diamond blade on a JEOL LTD JUM-7 ultramicrotome, contrasted with uranyl acetate and lead citrate before being observed under a Zeiss LEO 906E transmission electron microscope.

For EVs, 5µL of PBS containing EVs were fixed in 2% formaldehyde for 2h and then placed onto 200-mesh nickel grids coated with Formvar. To prevent non-specific binding, the grids were incubated in 5% BSA for 10 min. Monoclonal primary antibody CD63 (1µg/mL, Santa Cruz Biotechnology) was added for 1h. Protein A conjugated with colloidal gold (1/30, Electron Microscopy Sciences) was added for 30 min. For counterstaining, the EVs were exposed to uranyl-oxalate, followed by immersion in methylcellulose-AU for 10 min. After air-drying, they were visualized via TEM, digitally recorded and quantified.

### EVs-induced cell proliferation

EVs were isolated from two sources: HPSC controls and HPSC stimulated by 20µg/mL OxLDL1 for 24h. These EVs were normalized according to the number of secreting cells (250 000), resuspended in 50µL of PBS and used as stimuli in experiments conducted in a 24-well plates over 24h. Experiments involved both PC3 cell line epithelial cells and HPSC, cultured according to standard protocols^28,29^. Notably, primary cultures from which the EVs were extracted served as both EVs sources and recipient cells, enabling an analysis of EV effects on stromal cells from the same individual. The impact of EVs on cell proliferation was also examined through Ki-67 immunocytochemistry as detailed previuosly.

### Statistical analysis

Student’s t-test and analysis of variance (ANOVA) were conducted, followed by Tukey’s post hoc comparisons when necessary. To denote statistically significant differences between means, the following reference system was adopted: (*) for p < 0.05, (**) for p < 0.01, or (***) for p<0.001. GraphPad Prism 9.4 software was employed for performing the statistical tests and generating graphical representations of the data. All experiments were performed at least three times, each in triplicate.

## 5 Results

### Mice on HFD dysplayed an increase in prostate cell proliferation

Immunohistochemistry of the proliferative marker Ki-67, performed on ventral prostate sections, revealed a significant 2-fold increase in the number of positive cells in mice fed on HFD compared to those in the control group. Both stromal and epithelial compartments showed increments in prostatic cell proliferation, which was positively correlated with circulating total cholesterol levels mice (Fig. 1). These findings support the hypothesis that pro-atherogenic conditions have a detrimental effect on the prostate, promoting cellular proliferation.

**Figure 1:**
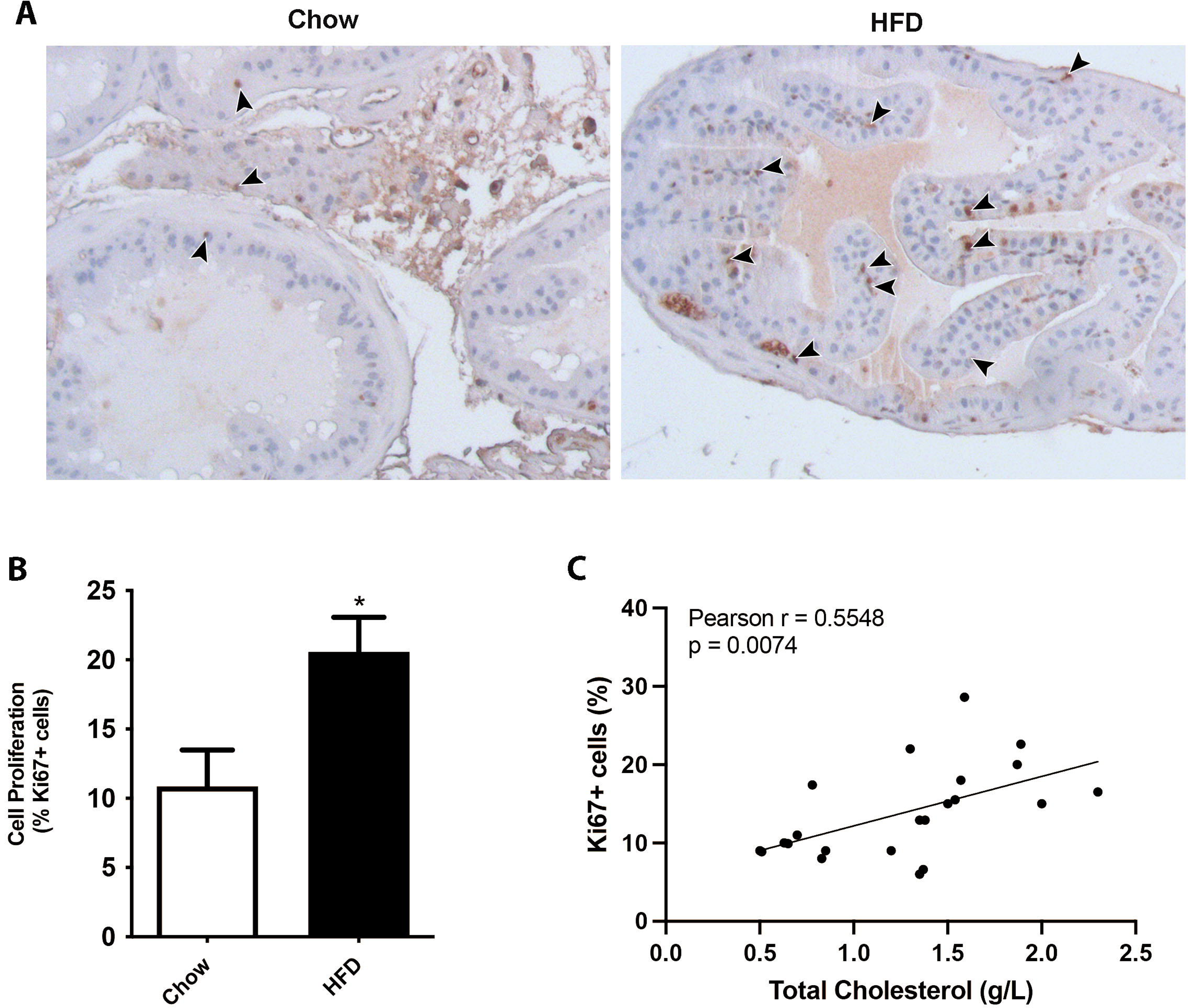
High-fat diet-induced hypercholesterolemia promotes cell proliferation in the prostate gland. Mice were fed on chow or high-fat diet (HFD) for 12 weeks and their prostates processed and analyzed by Ki-67 immunohistochemistry. A) Representative images of prostate glands immunostained for the proliferation marker Ki-67 showing an increase in HFD fed mice (brown nuclei, arrowheads). B) Quantification of Ki-67 positive cells in ventral prostates from chow or HFD fed mice (mean ± SEM; n =10 per group; *p < 0.05). C) Correlative analysis showing serum cholesterol levels and prostatic Ki67 positive counts (n=20, Pearson’s correlation test).

### Metformin inhibts OxLDL-induced cell proliferation in HPSC

Primary cultures identified as HPSC-1, HPSC-2, HPSC-3, HPSC-4, HPSC-5, HPSC-6, HPSC-7, and HPSC-8 were established from 8 BPH patients and stimulated with OxLDL. Following 15 days of growth, HPSC exhibited a spindle-shaped morphology and a well-developed cytoskeleton; ultrastructural and histochemical analyses demonstrated a myfibroblastic profile in these primary HPSC (Supplementary Figure 1).

The effects of OxLDL1 and OxLDL2 on HPSC were compared at various concentrations. As shown in Fig.2A, 5µg/mL did not induce significant changes in cell viability, apoptosis, and proliferation compared to controls. 100µg/mL OxLDL exhibited substantial toxicity for both OxLDL types, resulting in low cell viability and reduced proliferation. When stimulated with a 20µg/mL concentration, HPSC showed significant changes in cell proliferation in response to both OxLDL1 and OxLDL2 compared to controls. However, this concentration also led to elevated cell death for OxLDL2 (Fig.2A). Based on these findings, 20µg/mL of OxLDL1, the moderately oxidized form, was selected for all subsequent experiments. Accordingly, HPSC stimulated by OxLDL1 for 24h exhibited an increase in cell viability and cell proliferation, as determined by resarzurin absorbance and BrdU assay (Fig.2B) as well as by Ki-67 immunocytochemistry (Fig.2C) in the 8 primary cultures. To check the specificity of these effects, we treated the monocytic THP-1 cell line with OxLDL1 and OxLDL2 at a concentration of 20µg/mL for 24h. Despite being taken-up by THP-1 cells (Supplementary Figure 2), OxLDL did not promote significant changes in cell viability (Fig.2D) or proliferation (Fig.2E).

**Figure 2:**
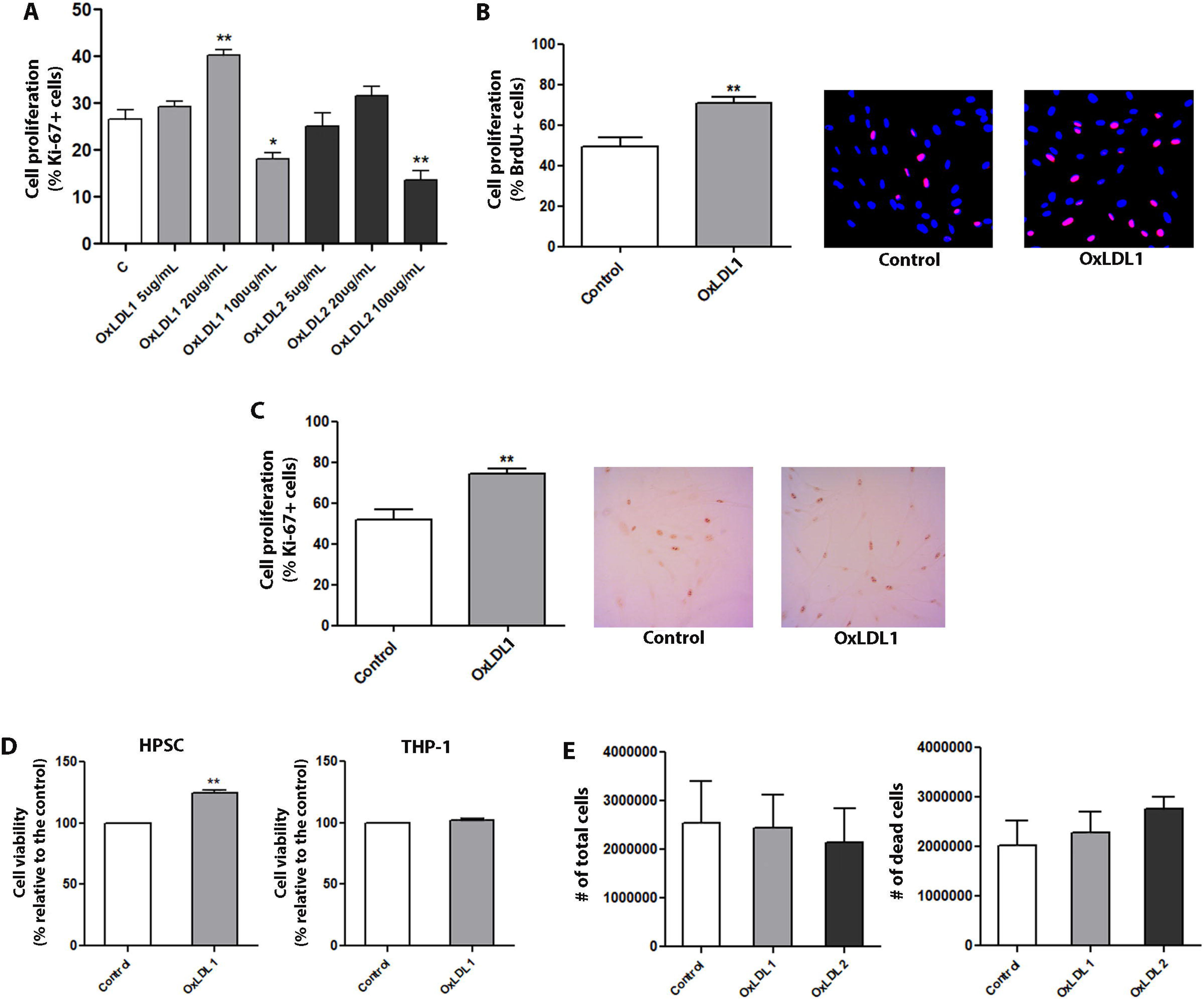
OxLDL at 20µg/mL increases cell proliferation and viability of HPSC. A) HPSC from patient HPSC-1 were used for the initial proliferation assay. Ki-67 immunocytochemistry was performed to evaluate the effects of two types of OxLDL: moderately oxidized (OxLDL1) and highly oxidized (OxLDL2). Although OxLDL1 at 20 and 100µg/mL and OxLDL2 at 100µg/mL showed changes in cell proliferation, OxLDL at 20µg/mL was the only stimulus without signs of cell toxicity. Additionally, OxLDL at 20µg/mL induced an increase in cell proliferation (**p < 0.01 vs. control). B) Quantification and representative merged images of DAPI/BrdU showing the proliferative effect of OxLDL1 at 20 µg/mL for 24 hours vs. control (**p < 0.01), evaluated in different primary cultures of HPSC by BrdU incorporation (n=7). C) The same proliferative effect of OxLDL1 at 20µg/mL for 24 hours vs. control (**p < 0.01) was evaluated in primary cultures of HPSC by Ki-67 assay (n=8). D) Resazurin absorbance assay for HPSC and THP-1 stimulated with OxLDL1 at 20µg/mL for 24 hours was carried out to assess cell viability. OxLDL1 at 20µg/mL increased cell viability compared to control (**p < 0.01). E) Assays with OxLDL1 and OxLDL2 stimuli in THP-1 cells evaluating the number of total live and apoptotic cells. Neither OxLDL1 nor OxLDL2 exerted changes in the cell number of THP-1 cells. All tests were performed in replicates, each in triplicate. Error bars represent mean ± SEM.

Treatment of HPSC with metformine at both doses (2 and 10 mM) abrogated OxLDL-induced cell proliferation and viability, as determined by BrdU incorporation, Ki-67 immunocytochemistry, total cell count, and Resazurin assays (Fig.3). Co-stimulation with atorvastatin did not result in a reduction in OxLDL-induced cell proliferation. Atorvastatin alone did not exhibit inhibitory effects on cell proliferation. This finding was consistent across all primary cultures evaluated, which suggests a potential secondary therapeutic role for metformin, beyond its hypoglycemic effects, in attenuating the proliferative processes in BPH under dyslipidemic conditions.

**Figure 3:**
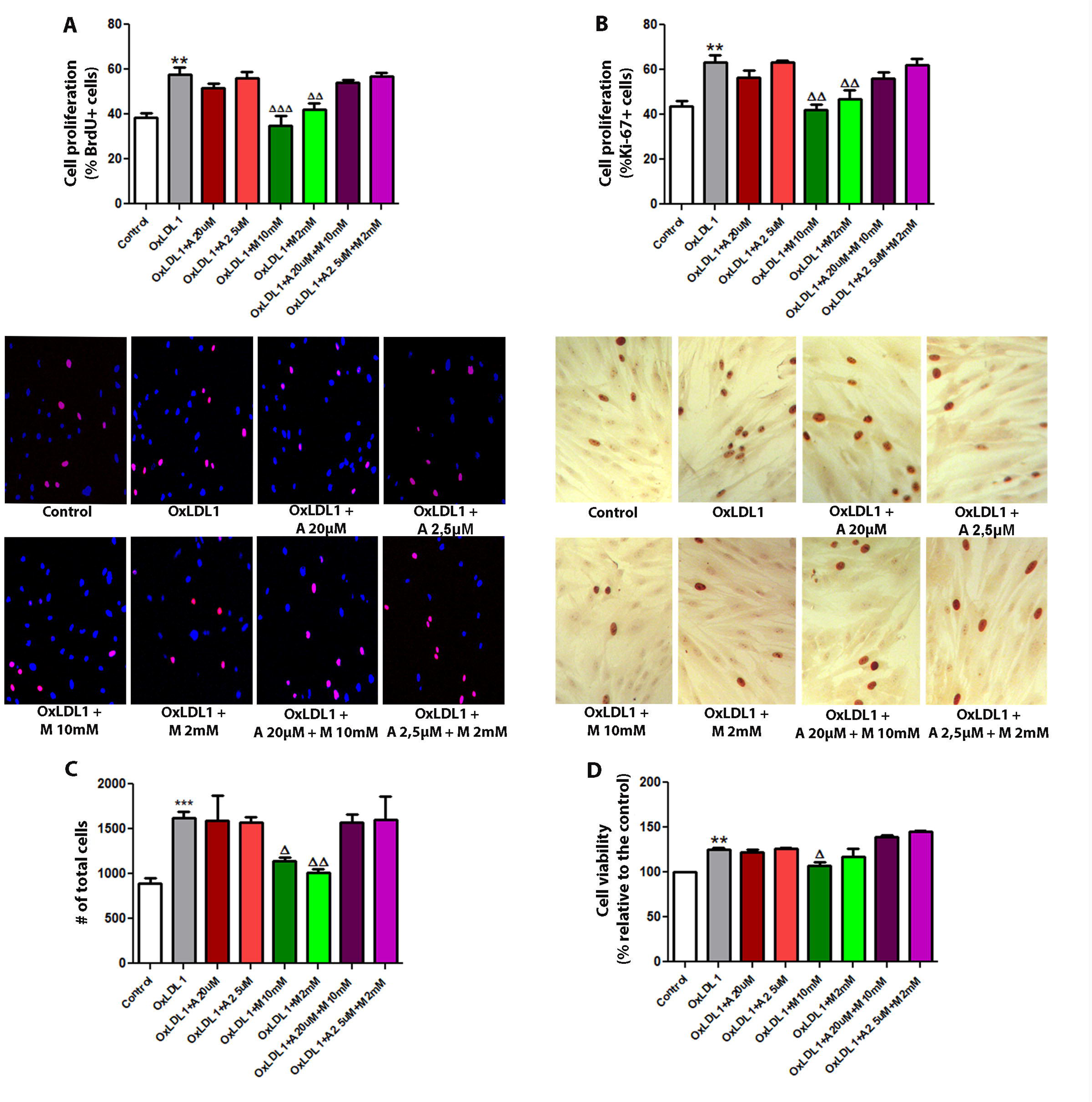
Metformin inhibits the proliferative effect of OxLDL on HPSC. Cells were stimulated with 20µg/mL of OxLDL1, and atorvastatin, metformin, or a combination of both were used as co-stimuli, each at low and high doses for 24 hours. A) Cell proliferation assays by BrdU incorporation were carried out in primary cultures of HPSC. Images of cells are representative of patient HPSC-6. OxLDL1 enhanced cell proliferation compared to the control (**p < 0.01), while metformin inhibited the proliferative effect induced by OxLDL1 (**p < 0.01 and ***p < 0.001 vs. OxLDL1). B) The same effect was observed by Ki-67 immunocytochemistry on cells from the same patient (**p < 0.01 for OxLDL1 vs C, and **p < 0.01 for metformin vs OxLDL1). C) Determination of the total cell number performed on cells from patient HPSC-5. OxLDL1 increased the cell number compared to the control (***p < 0.001), while metformin reduced the cell number compared to OxLDL1 (*p < 0.05 and **p < 0.01). D) Cell viability assay evaluated by resazurin absorbance was performed on cells from patient HPSC-5. All the tests were performed in two replicates, each in triplicate. Bars represent mean ± SEM.

### OxLDL induced morphological changes in HPSC

HPSC exhibited a myofibroblastic phenotype, including a prominent rough endoplasmic reticulum, Golgi apparatus, secretion granules, and peripherally located microfilament bundles. OxLDL treatment resulted in markly ultrastructural alterations (Fig.4A), along with an increase in the amount of organelles associated to cellular metabolism and contractile activity (Fig.4B). Signs of cellular stimulation after OxLDL included the ocurrence numerous nucleoli, proteinopoietic organelles, myofilaments with focal densities, and scarce fibronexuses. External fibronectin fibril were rarely observed. These findings align with increased cell proliferation, indicating that OxLDL promotes a more agresive morphological phenotype in HPSC.

**Figure 4:**
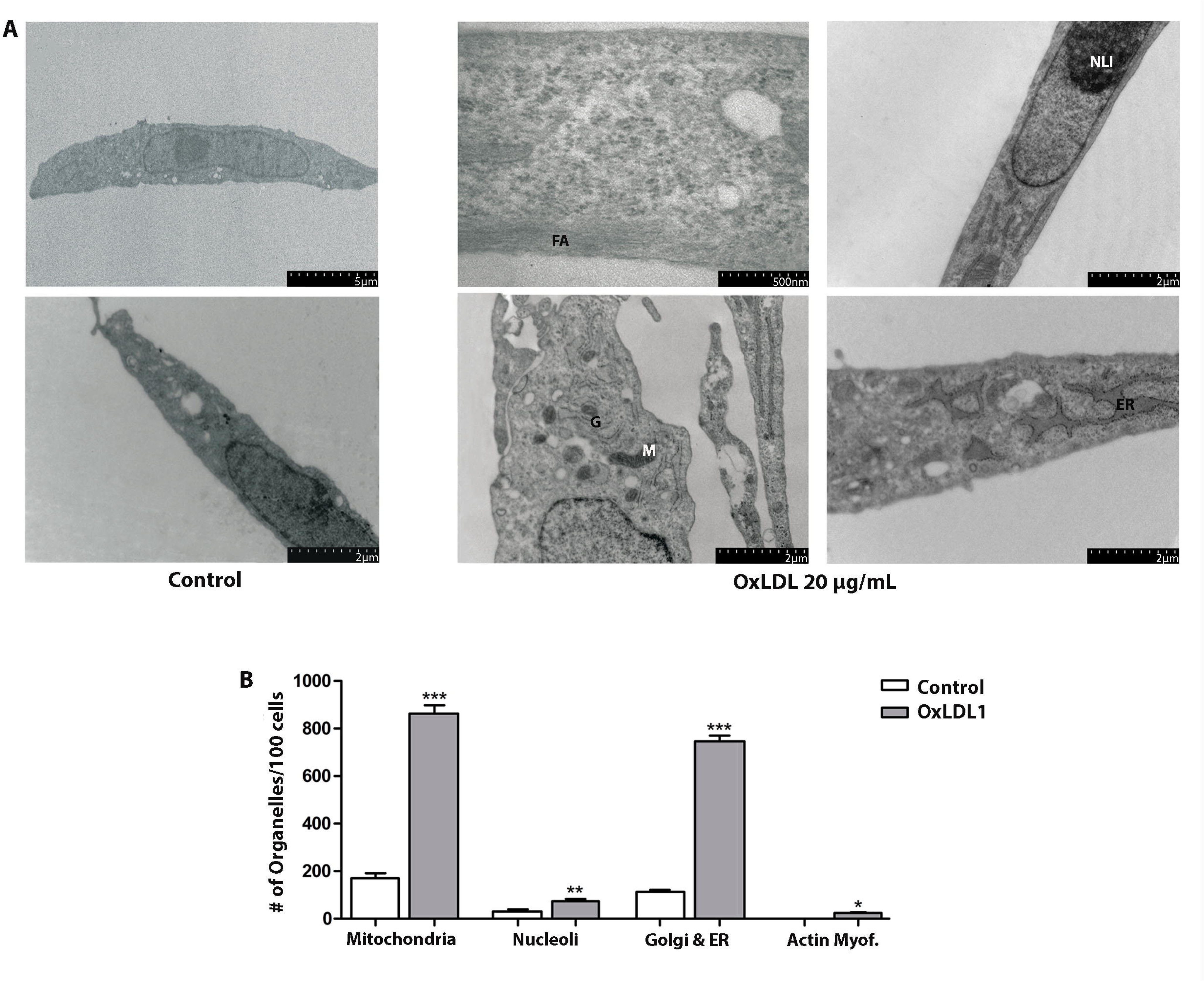
OxLDL causes morphological and ultrastructural changes in HPSC. A) Ultrastructure of HPSC derived from patient HPSC-1, treated with OxLDL1 at 20µg/mL for 24 hours, and evaluated by TEM. Cells stimulated by OxLDL displays frequent actin filaments (FA), dilated endoplasmic reticulum (ER), numerous mitochondria (M), Golgi apparatus (G), and prominent nucleoli (NLI). B) Quantification of organelles observed by TEM for HPSC from patient HPSC-1, control vs. treated with OxLDL1 at 20µg/mL for 24 hours. There was a significant increase in the number of mitochondria (***p < 0.001), nucleoli (**p < 0.01), Golgi and endoplasmic reticulum (***p < 0.001), and actin myofibrils (*p < 0.05) when stimulated by OxLDL1 compared to the control. All tests were performed in two replicates, each in triplicate. Bars represent mean ± SEM.

### EVs derived from OxLDL-treated HPSC showed proliferative activity

The electron microscopy analysis of EVs isolated by ultracentrifugation from control and treated HPSC revealed mainly concave, calyx-shaped (Fig.5A), with a peak in the range of 15-30 nm-sized structures (small EVs, previously recognized as exosomes). Immunogold staining for the tetraspanin CD63 also confirmed the identity as EVs (Fig.5B). OxLDL induced a significant increase in smaller EVs, particularly in the 15 to 60 nm range (Fig.5C).

**Figure 5:**
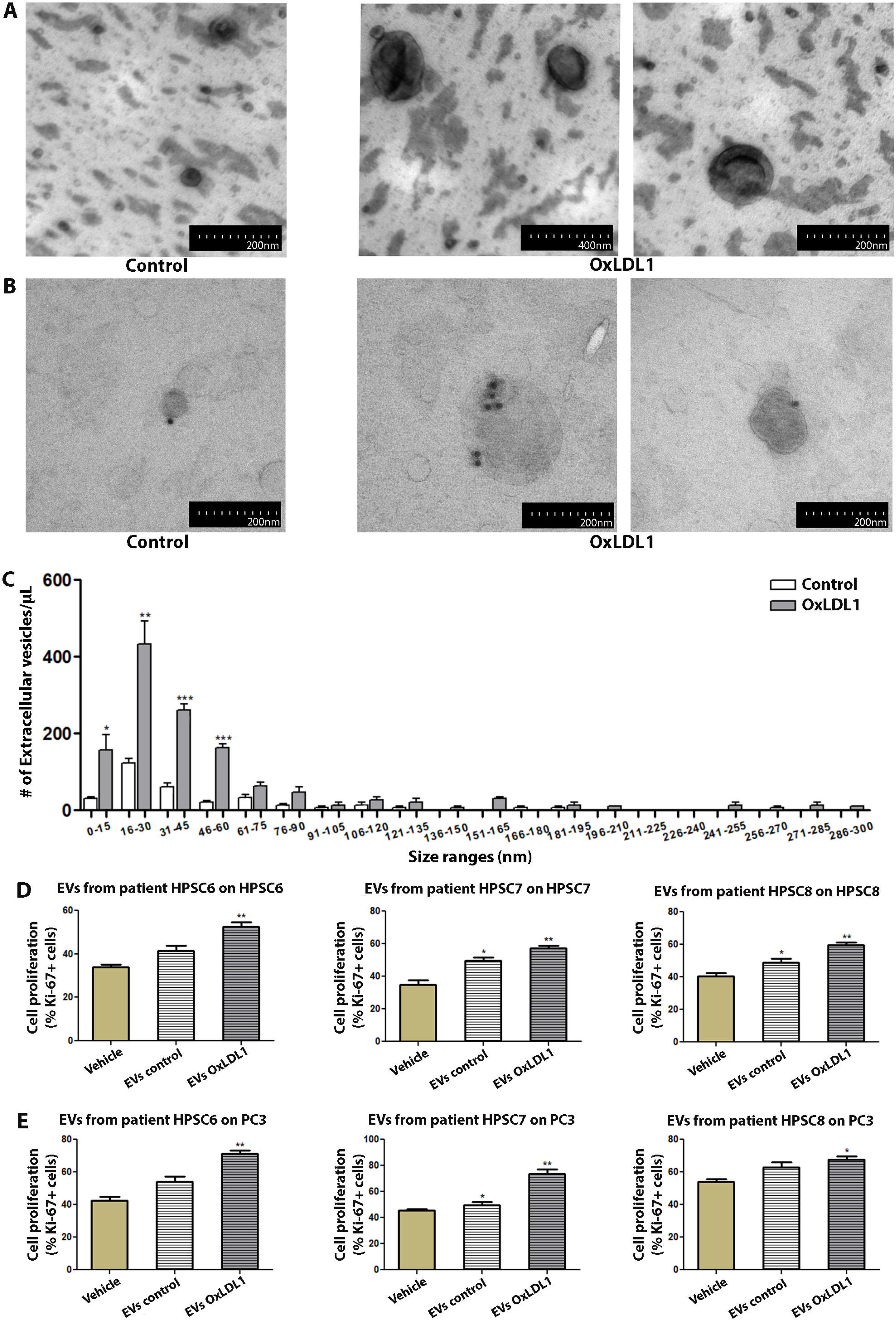
EVs derived from OxLDL-stimulated HPSC exert a pro-proliferative effect on prostatic cells. A) Images of HPSC-derived EVs from patient HPSC-4, treated with OxLDL1 at 20µg/mL for 24 hours, isolated by ultracentrifugation and visualized by TEM. B) Immunolabeling for CD63 in EVs derived from patient HPSC-4, using colloidal gold particles. C) The histogram of size range distribution of EVs derived from HPSC of patient HPSC-4 shows an increase in the number of EVs, mainly in the lower ranges. EVs obtained from HPSC stimulated with OxLDL1 at 20µg/mL for 24 hours (EVs OxLDL1) and control (EVs control) were used as stimuli on HPSC from the same patient (D) and on prostatic epithelial cells from the PC3 line (E). EVs from OxLDL-stimulated HPSC induced a significant proliferative effect on HPSC and PC3 cells in all assays (**p < 0.01 and *p < 0.05 vs. vehicle). The proliferative effect was evaluated by Ki-67 and all tests were performed in three replicates, each in triplicate. Bars represent mean ± SEM.

Additionally, EVs from supernatants of vehicle- and OxLDL-treated cells from three different BPH patients were isolated and used to stimulate HPSC and PC3 cells (Fig.5D-E). OxLDL-induced EVs significantly increased cell proliferation compared to vehicle- or EVs control-treated HPSC. Interestingly, OxLDL-pulsed EVs showed similar proliferative activity on the PC3 epithelial cell line, suggesting that EVs can promote cell proliferation in autocrine as well as paracrine manner in the context of BPH.

## 6 Discussion

The association between dyslipidemia and abnormal prostatic growth has been extensively investigated in the last decade. Although compeling evidence exists from epidemiological studies, limited information is still available on the cellular mechanisms explaining such correlation. Our study found that HFD, which elevates LDL levels, induces prostatic cell proliferation *in vivo*. Additionaly, primay cultures of HPSC from patients with BPH showed increased proliferation in response to OxLDL, which was inhibited by metformin. Furthermore, OxLDL augmented the secretion of pro-proliferative EVs by HPSC, suggesting a potential mechanism underlying BPH progression in pro-atherogenic and dyslipidemic conditions.

LDL and its oxidized forms are known to cause vascular and microcirculatory lesions in various tissues in the contexts of dyslipemia, MS, chronic inflammation, and hormonal imbalance^14,30^. *In vitro,* OxLDL can induce proliferation on human endothelial cells^31,32^ and on prostatic PC3 and LNCaP cell lines^33^, and it has been implicated in promoting angiogenesis in C4-2 cells through LOX-1 receptor activation^34^. Moreover, Haga *et al.* have linked OxLDL to prostatic growth, attributing it to atherosclerosis-induced chronic inflammation via LOX-1^35^. Our ultrastructural observations suggest an increment in both metabolic function and contractile activity of HPSC after OxLDL, which promotes features of aggressive myofibroblasts^36,37^. Similar changes have been reported in cardiac myofibroblasts exposed to OxLDL, exacerbating a pro-fibrotic phenotype^38–40^. Our findings underscore the pathological role of OxLDL in BPH and support the hypothesis linking dyslipidemia, inflammation and MS components with benign prostatic growth and LUTS^14^.

Metformin and atorvastatin are widely prescribed for pro-atherogenic conditions like dyslipidemia, diabetes, and MS^41^. Notably, these drugs have also shown potential in regulating cell proliferation in different contexts^42–44^. Our results demonstrated antiproliferative effects for metformine, while atorvastatin did not impact OxLDL-induced proliferation of HPSC *in vitro*. However, the combination of metformin and atorvastatin failed to counteract cell proliferation, indicating intricate signaling pathways involved. Different anti-proliferative mechanisms of action for metformin have been reported, including mTOR inhibition, AMPK activation, modulation of protein synthesis and cell growth, and the activation of p53^45^. This discovery offers insight into metformin effects and a rationale for potential application in patients with BPH.

Over the last years, studies have demonstrated that dyslipidemic states can increase the secretion of EVs by various cell types, including macrophages^21^, endothelial cells^20^, and platelets^22^. Furthermore, OxLDL-induced EVs secretion appears to be mediated by the activation of scavenger receptors LOX-1 and CD-36^46^. Here, we demostrate for the first time that OxLDL estimulates the production and secretion of pro-proliferative EVs by HPSC from patients with BPH. Previous articles have shown that tumoral prostatic cells and the cell line WPMY-1 secrete EVs^47–49^. However, it remains uncertain whether these EVs share morphological and functional features with classical prostasomes or other microparticles secreted by the prostate gland^48^. These findings highlight the prostate stromal cells capacity to secrete EVs into and modulate the extracellular milieu. In fact, the interactions between stromal and epithelial cells control the strict balance in pro- and anti-proliferative signals, which is dysregulated in BPH. In this scenario, autocrine and stroma-to-epithelium communications mediated by EVs appear to be an important player in the proliferative proccess promoted by dyslipedimia. This effect can be ascribed to the multiple EV cargoes; for instance, miR-92a-3p, which inhibits the expression of thrombospondin-1, has been shown to regulate EV-induced cellular proliferation in the context of OxLDL challenge^49^.

## 7 Conclusions

In summary, our study provides evidence that the dyslipidemic environment is associated with cell proliferation in the prostate gland and that OxLDL increases proliferation and metabolic activity in human stromal cells from BPH patients. Additionally, the proliferative activity relies on the secretion of EVs by HPSC, facilitating cellular communication between the stromal and epithelial compartments of the prostate gland.

## 8 Acknowledgements

We would like to thank Carolina Leimgruber and Lucia Artino for TEM imaging, and Sofía Rossetto for her excellent technical assistance.

## Ethics approval statement

This study adhered to the NIH Guidelines. The work involving animal samples was approved by the local ethical committee (CICUAL, UNC, Argentina). Additionally, the work involving human samples was approved by the Bioethics Committee of Sanatorio Allende (Córdoba, Argentina).

## Supplementary

**Supplementary Figure 1: Morphology of HPSC from primary cultures derived from patients with BPH.** Cellular morphology of HPSC shows their characteristic spindle-shaped morphology with the nucleus located in the central zone: A) HPSC stained with Toluidine Blue. B) HPSC visualized using a fluorescence microscope with Phalloidin labeling for cytoplasmic actin filaments (red) and DAPI for the cell nucleus (blue). C) HPSC imaged with differential interference contrast (DIC) microscopy. D) HPSC observed by transmission electron microscopy (TEM).

**Supplementary Figure 2: OxLDL is internalized by THP-1 cells, but it does not induce changes in cell morphology or ultrastructure.** A) Internalization of OxLDL molecules (yellow arrows) by THP-1 cells was observed by TEM. B) The ultrastructure of THP-1 cells was evaluated by TEM. No changes were observed when treating cells with OxLDL1 at 20µg/mL for 24 hours. Scale bars: 5µm. C) The organelle count for THP-1 cells, comparing control vs. OxLDL1 treatment, did not show statistically significant differences for *p < 0.05. The test was performed in two replicates, each in triplicate.

