## Supplementary figures and images for "Extracellular vesicles mediate OxLDL-induced stromal cell proliferation in Benign Prostatic Hyperplasia"

### Supplementary Fig 1

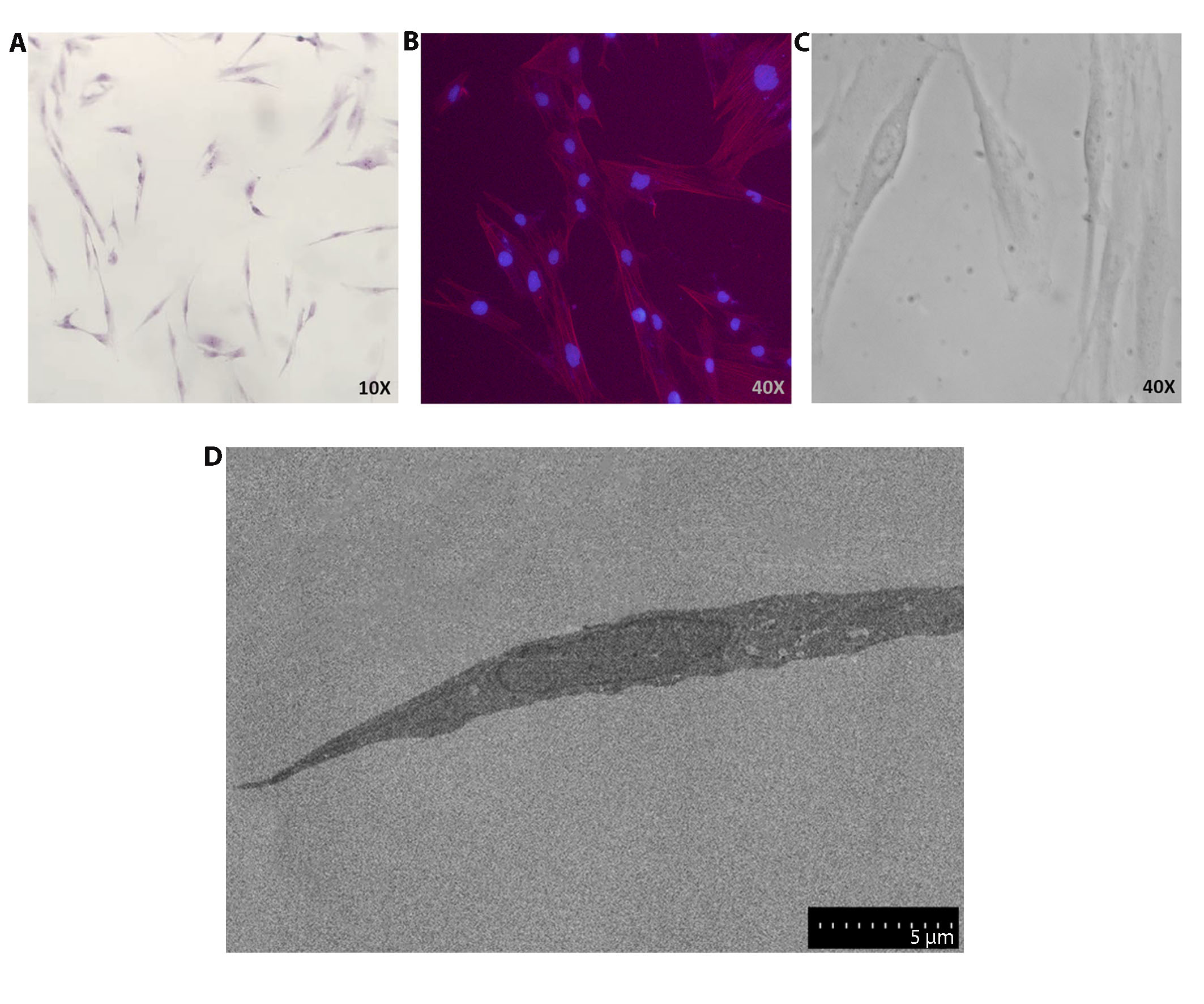

### Supplementary Fig 2

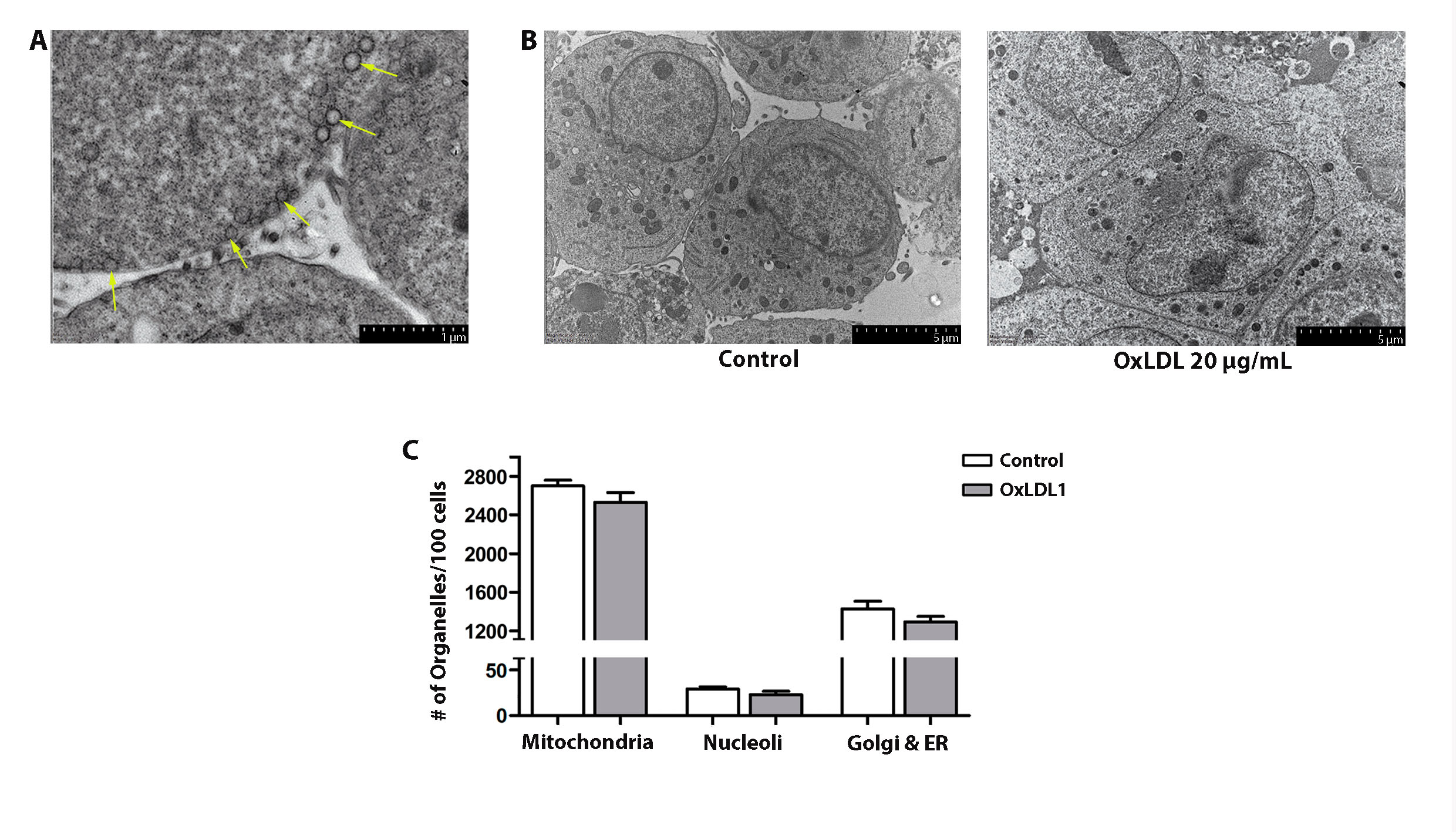
